# A new trait in a well-studied ant: low aggression between workers from *Lasius niger* (Hymenoptera, Formicidae) nest complexes

**DOI:** 10.1101/2024.07.23.604725

**Authors:** Stanislav Stukalyuk

## Abstract

The formicine ant *Lasius niger* is one of the most abundant and most intensively studied Palaearctic ant species and was believed so far to form exclusively monogynous colonies, to spread via single-queen dispersal and to found claustrally. Two closely neighboring nest complexes of *Lasius niger* were observed in an abandoned field near Kyiv / Ukraine in 2017-2020. Nest complex A includes 14650 mounds on an area of 11.8 ha and nest complex B 15600 mounds on 13.3 ha. The nests within the nest complexes are connected via a developed trail network. Aggression tests revealed a low level of aggression between workers from different zones of the same nest complex but increased aggression in confrontations of nest complex workers with workers from a remote monodomous population. Reduced aggression between workers facilitate the integrity of the nest complex and its rapid growth.

## Introduction

Polydomy in ants is a form of social organization in which multiple physically separated nests are socially connected by the exchange of workers, brood and food. The alternative condition is called monodomy in which the whole ant colony is housed in a single nest and behaves aggressively against other conspecific monodomous colonies. Polydomous systems occur more rarely in colonies with a single reproductive queen (monogyny) and more frequently in those with multiple queens (polygyny) (e.g. Debout et al. 2007; Seifert 2018). Depending on the species’ potential and environmental conditions, polydomous colonies can reach an enormous population size and cover a large area. The extreme expression of this life form is called a nest complex and was defined by Seifert (2018) as “a large to huge polydomous-polygynous colony with permanent connections between the nests and workers not showing mutual aggression even if originating from nests kilometers apart”.

Supercoloniality is a typical trait of many invasive (neozootic) ant species and is one reason for their success in the newly colonized territories. In Europe north of the Alps ten invasive ants species have occupied urban and rural outdoor habitats and nine of these are supercolonial and became notorious pest species (Seifert 2020a, b). In extreme cases, nest complexes of invasive ants contain millions of nests (Giraud et al. 2002; Sunamura et al. 2009; Van Wilgenburg et al. 2010; Moffet 2012). The number of nests can range from thousands in *Crematogaster subdentata* Mayr, 1877 (Stukalyuk and Netsvetov 2018) to millions in *Linepithema humile* (Mayr, 1868) or the invasive garden ant *Lasius neglectus* Van Loon et al. 1990 (Tartally 2006, reviewed in Seifert 2018).

Nest complexes of non-invasive, autochthonous ants can also reach impressive size. Examples of very big nest complexes in the temperate to subboreal zones of the Palaearctic are found in particular in the ant genus *Formica*: *Formica yessensis* Wheeler, W.M., 1913 in Japan with 45000 nests on an area of 270 ha (Higashi and Yamauchi 1979), *Formica exsecta* Nylander, 1846 in Romania with 3347 nests on 21.8 ha (Markó et al. 2012), *Formica foreli* Bondroit, 1918 in Germany with 2550 nests on 6.2 ha (Bönsel and Busch 2003) and *Formica paralugubris* Seifert, 1996 in Switzerland with 1200 nests on 70 ha (Cherix and Bourne 1980).

Ants of the genus *Lasius* are considered to be rarely polygynous and thus the likelihood to form polydomous structures is low. Among the 56 Palaearctic species of subgenus *Lasius* s. str. (Seifert 2020c) four facultatively to obligatory polygynous species are known so far. *Lasius austriacus* Schlick-Steiner 2003 is weakly polygynous or oligogynous and forms polydomous subterranean colonies (Pedersen and Cremer 2008). *L. sakagamii* Yamauchi & Hayashida 1970 and *Lasius neglectus* are highly polygynous and may form true nest complexes (Yamauchi et al. 1981; Espadaler et al. 2004; Tartally 2006; Stukalyuk and Radchenko 2018). *Lasius precursor* Seifert 2020 is moderately polygynous and polydomous (Cremer et al. 2008) and was considered by Seifert (2020c) as a model for transition from a largely monogynous-monodomous social type (exemplified by the sister species *Lasius turcicus* Santschi 1921) to a supercolonial type (exemplified by the closely related species *L. neglectus*).

*Lasius niger* (Linnaeus, 1758), as the most abundant and best-studied member of the subgenus *Lasius* s. str., shows a continuous distribution from West Europe (10°W) to the eastern Baikal region (108°E). It is by natural distribution mainly an element of the northern steppe zone and the transition zone from steppe to temperate forest but following the spread of human culture there was a strong range expansion into rural and urban habitats even of the south boreal zone (Seifert 2018, 2020c). Plenty of studies on diverse aspects of biology have been conducted in this ant. Swarming flight and mating scenarios were intensively studied by Boomsma and Leusink (1981), Boomsma and van der Have (1998), Fjerdingstad et al. (2002) and Van der Have et al. (2011). Colony foundation and colony demography were investigated by Mrazek (1906), Eidmann (1928), Goetsch (1938), Sommer and Hölldobler (1992, 1995), Aron and Passera (1999) and Aron et al. (2009). The observations of these authors resulted in the following consistent picture: after mass swarming and aerial mating *Lasius niger* gynes perform claustral colony foundation either alone (in haplometrosis) or in temporary cooperation of several gynes (in pleometrosis). The last mentioned seven authors agree in stating that pleometroses are terminated by mortal fighting between foundresses after the first workers have eclosed from pupae resulting in the survival of only one queen. The most fertile queen is more likely to be fed by the workers and thus to survive the fight (Sommer and Hölldobler 1992, 1995; Aron et al. 2009). As result, a monodomous-monogynous colony structure is established with workers of different nests behaving mutually aggressive. This presented the generally accepted view on this ant so far.

In this paper, we describe for the first time the spatial structure, connectivity and intraspecific behavior in a huge nest aggregation of *Lasius niger* discovered near Kyiv /Ukraine. We argue that this nest aggregation it meets the following criteria: (1) nests are interconnected by a network of above-ground trails and subterranean tunnels within which workers move freely, (2) workers from close to remote parts of the colony show no or only little aggression to each other.

## Materials and Methods

### Study area

Two closely-neighboring, large nest complexes of *L. niger* were discovered near the city of Vyshneve (Kyiv-Sviatoshyn Region, Kyiv District) in Ukraine (50.3906°N, 30.3364°E, 179 m a.s.l.). Including patches with no mounds detected, the nest complexes cover an area of 27 ha. The site is a flat terrain with small elevations, dry and clayish soil and has been arable land in the past. Cultivation has been gradually abandoned since 2010. Today, the area is dominated by segetal, semi-ruderal and ruderal communities of annual, biennial, and perennial plants, namely grass species from classes *Stellarietea mediae* and *Artemisietea vulgaris* (Mosyakin and Fedoronchuk 1999) as well as remnants of the former crop communities (cereals). The vegetation has a mean height of 20 to 30 cm and a surface coverage of 40 to 65%. There are also spots where the grass stand reaches a height of 50 to 70 cm.

### Observation and recording methods

Field studies were conducted in 2017 to 2018. To check if this huge complex of *L. niger* nests could represent a true nest complex, we investigated the boundaries of the complex, the network of above-ground trails and subterranean tunnels between, the level of aggression between ants in different zones of the complex and the behavior of gynes in pleometrosis experiments which allows conclusions on the likelihood of coexistence of several queens in the same nest mound (polygyny).

Assessing density and size of nest mounds was done according to the methods of Dlussky (1965) and Zakharov and Goryunov (2009) by walking along 10-m-wide line transects covering the whole area. This was done in April 2017 when the vegetation cover was low and the mounds were clearly visible (Figure 1).

**Fig. 1.**
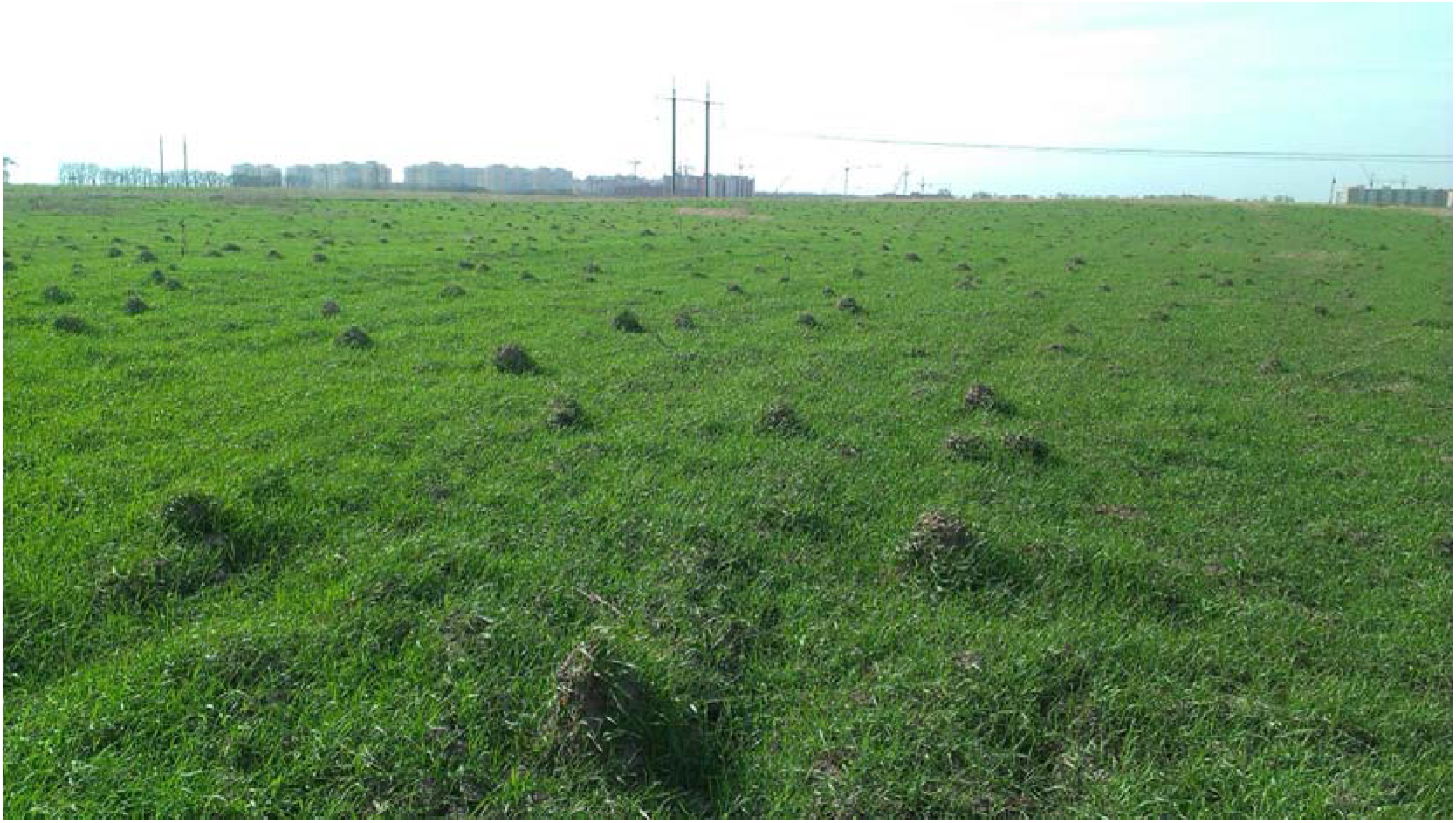
The *Lasius niger* nest complex B, sectors 7, 8.

Additionally, height and diameter of nest mounds were measured. A total of 118 transects were allocated with lengths differing in dependency from the shape of the two nest complex territories. Based on transect data, a mosaic of 23 sectors was established which contain nest mounds from different density classes: Class I with 1 to 5 mounds/100 m^2^, Class II with 6 to 10 mounds/100 m^2^, Class III with 11 to 15 mounds/100 m^2^, and Class IV with ≥16 mounds/100 m^2^. Locations of populated sectors and of areas with no mounds were mapped by placing their geographic coordinates on Google Maps. The distinction of two separate nest complexes A and B was based on their complete isolation by unpopulated arable land and on aggression tests. In three 100 m^2^ quadrats, detailed mapping of the following features was performed: major nests with clearly visible mounds, underground nests not covered by mounds, auxiliary structures such as soil pavilions around the feeding sites and small, and above-ground trails and subterranean tunnels.

Twelve aggression tests between workers from mounds from different zones of nest complex B were conducted in late May 2018. Nests from the center of the nest complex were tested against nests distant from the center by 30, 250, 500 and 1000 meters, with three tests performed in each the four distance categories. In a control experiment, three tests between nests from the center of nest complex B and nests from a monodomous population located in a garden 2 km away (50.3957°N, 30.3449°E) were performed.

The test samples were collected by placing open test tubes with bait (freshly killed European June beetles *Amphimallon* sp.) next to a mound. The worker ants were allowed to feed on the bait until the tube was filled with around 100 workers. The tubes were then clogged with cotton wool and taken to the laboratory. This mode of sampling was considered to cause less stress than collecting the workers directly from the mound. In the pairwise confrontation tests in the laboratory, the tubes were tightly connected via their opened ends and the interaction between ants was videotaped during two minutes. This method requires much lower observation time and less technical expense as video recording has to monitor only a confined space instead of a large observation arena. However, this mode of encapsulated sudden confrontation is suspected to result in some aggression where under other circumstances no aggression would occur (see discussion). Similar to Wallis (1962), Batchelor and Brifa (2011), we classified the ants’ responses into five categories: swift thusts and drawbacks (1), threatening by mandible opening (2) or gaster flexion (3), seizing or clenching an enemy’s appendages (4) and attack with spraying of gaster secretions (5). Additionally, we counted the number of workers killed during confrontation. The aggressive behaviors of ants were scored from 1 to 4 with escalating intensity and then summed to a Total Agonistic Index (TAI). Swoops, bounces and threats were each scored 1, seizing was scored 2, attacks with apparent spraying of gaster secretions scored 3 and the number of killed ants scored 4. We calculated an index TAI_3_ considering only behavioral acts scored 1 to 3 and an index TAI_4_ additionally considering killed ants. Only observations of aggressive reactions are considered in this study whereas non-aggressive encounters such as friendly mutual antennation or trophallaxis were not recorded.

### Species determination

Ants were determined according to the key of Seifert (2018).

### Statistical analysis

We evaluated the Total Agonistic Index (TAI) data with a one-way ANOVA in comparing the TAI data of each of the five aggression test settings pairwise. Changes over time in the queen number in the pleometrosis experiments were evaluated by Fisher’s two-tailed exact test run with the software package R (R Development Core Team 2012). ANOVA tests and principal component analysis were performed with the software package SPSS 15.0.

## Results and discussion

### Species determination

Ants from both the nest complex are clearly determined as *Lasius niger*.

### Topography and size of the nest complexes

The examined nest complexes stretch over about 1200 m in southwest-northeast direction with a width ranging between 86 and 450 m (Fig. 2).

**Fig. 2.**
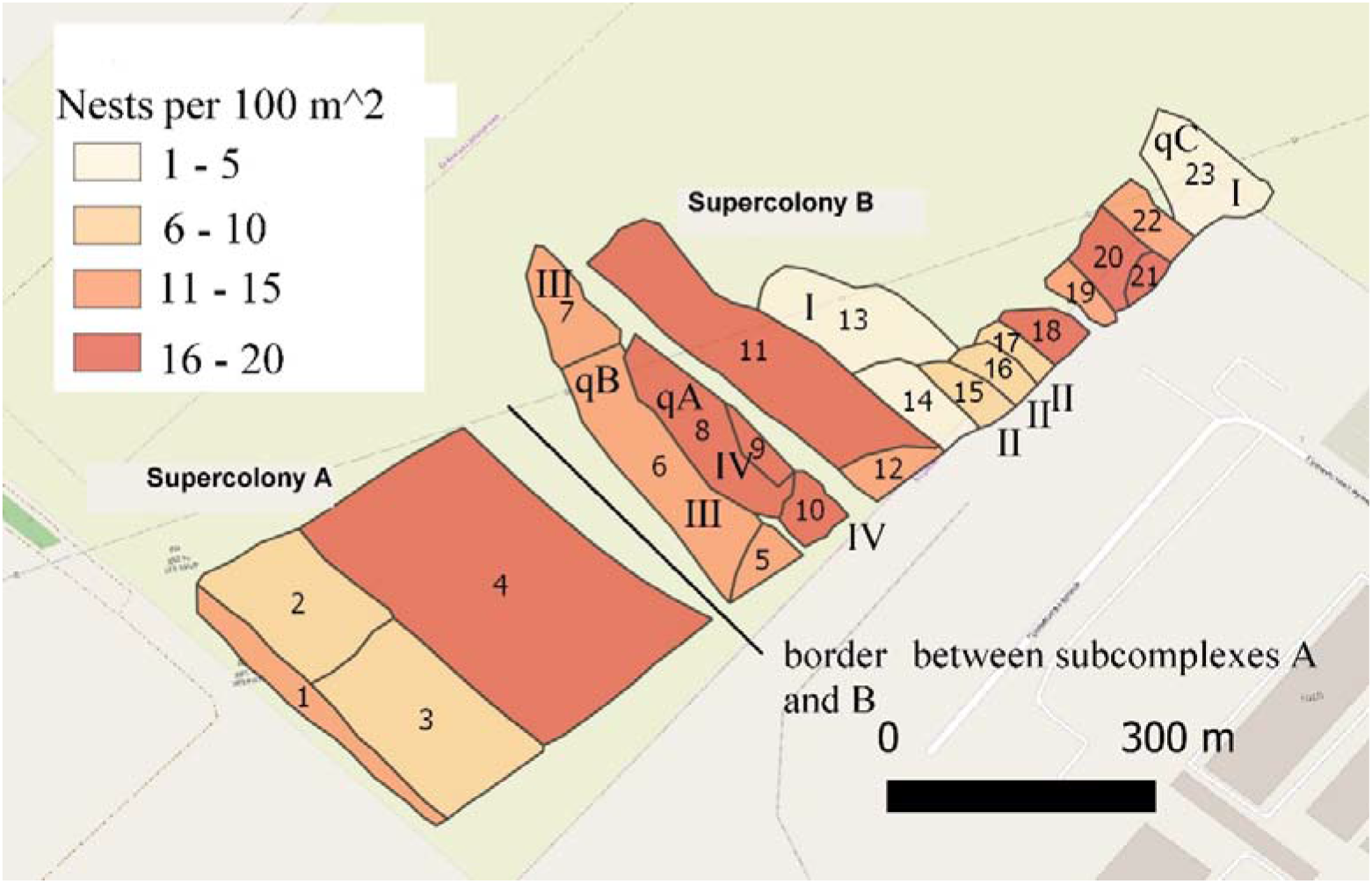
The map showing *Lasius niger* nest complexes A and B with sections of variable density.

There is a separation in two nest complexes by a wedge-shaped area of ploughed land of 20 to 120 m width. The southwestern nest complex A with sectors 1–4 is square-shaped, occupies an area of 11.8 ha and includes 14655 nests resulting in a nest density of 12.4 nests /100 m^2^. The corresponding data of the more elongated and irregularly shaped nest complex B with sectors 5–23 are 13.3 ha, 15599 nests and 11.7 nests / 100 m^2^. These nest density data are most probably an under-recording as the transect walks counted only the clearly visible mounds. As reported by Seifert (2017a) true nest densities are only recordable by carefully probing the whole ground. Seifert found a mean and maximum density of 17.6 and 108 nests / 100 m^2^ on 68 test plots in Central Europe belonging to a multitude of habitat types. Hence, the nest densities reported here do not provide an argument for supercoloniality. The same applies to the dimensions of the above-ground nest parts. According to the B. Seifert observations about the situation in Central Europe, the dimensions of mounds from the investigated nest complex, with a mean basal diameter of 35 cm and mean height of 24 cm (Tab. 1), correspond to the average situation in *Lasius niger*.

**Table 1.** Main characteristics of the *Lasius niger* nest complex B.

| Density class | Number of mounds per 100 m², $\bar{x} \pm \text{SE}$ | Average mound diameter, cm, $\bar{x} \pm \text{SE}$ | Average mound height, cm, $\bar{x} \pm \text{SE}$ |
| --- | --- | --- | --- |
| I | 15.9 ± 1.2 | 31.6 ± 0.6 | 25.6 ± 0.6 |
| II | 11.9 ± 1.1 | 39.0 ± 1.0 | 23.5 ± 0.7 |
| III | $11.5 \pm 0.7$ | $32.8 \pm 0.7$ | $20.2 \pm 0.7$ |
| IV | $4.4 \pm 1.0$ | $32.4 \pm 1.5$ | $21.7 \pm 4.7$ |
| Total | $11.1 \pm 0.8$ | $34.1 \pm 0.5$ | $22.9 \pm 0.4$ |

The spatial distribution of nests is also no conclusive argument for the presence of a nest complex. The two high-density zones in nest complex B (Fig. 2) might possibly indicate origins of colony extension by nest-fission. Yet, this is speculative as density and absolute size of nest mounds strongly depend from environmental factors and must not reflect the spreading history of a colony. As a consequence, we have to look for conclusive arguments for supercoloniality which are provided in the three next sections.

### Connections between the mounds

Fig. 3 shows the topography of above-ground trails and subterranean tunnels, connecting mounds within nest complex B.

**Fig. 3.**
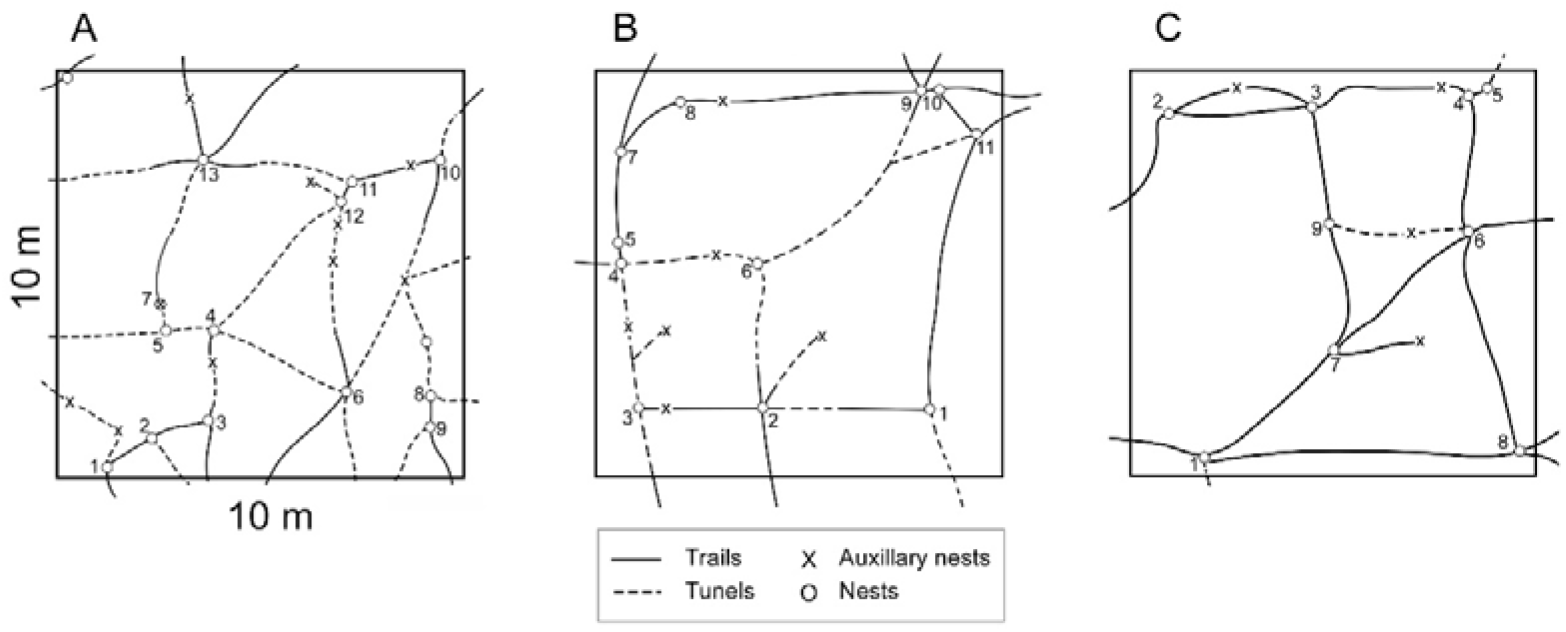
Nests and trails between nests in *Lasius niger* nest complex B. A – zone IV, B – zone III, C – zone I.

Direct observation of above-ground trails showed an intense traffic of workers. These multiple connections linking both major and auxiliary nests can be seen in different zones of the nest complex. Auxiliary nests are usually initiated by constructions of mineral soil covering aphid colonies, both subterranean or at base of plant stems, and may develop in the long term to major nests. Fig 4A shows an example of a trail network connecting 13 major mounds in a 100 m^2^ plot plus connections to seven auxiliary nests. A medium-sized *L. niger* nest with a basal mound diameter of 35 cm as found in our study (Tab. 1) is estimated to contain a population of 14 000 workers and the maximum population maintained in a monogynous nest seems to be approximately 60 000 (Boomsma et al. 1982). This maximum size is limited by the laying capacity of the single queen. The whole population of 13 major and 7 auxiliary nests mapped in the 100 m^2^ plot in Fig. 3A should be 250 000 workers at least. It appears most unlikely that such a big population can be maintained by a single queen.

### Aggression between workers

The results of the aggression tests are presented in Tab. 2.

**Table 2.**
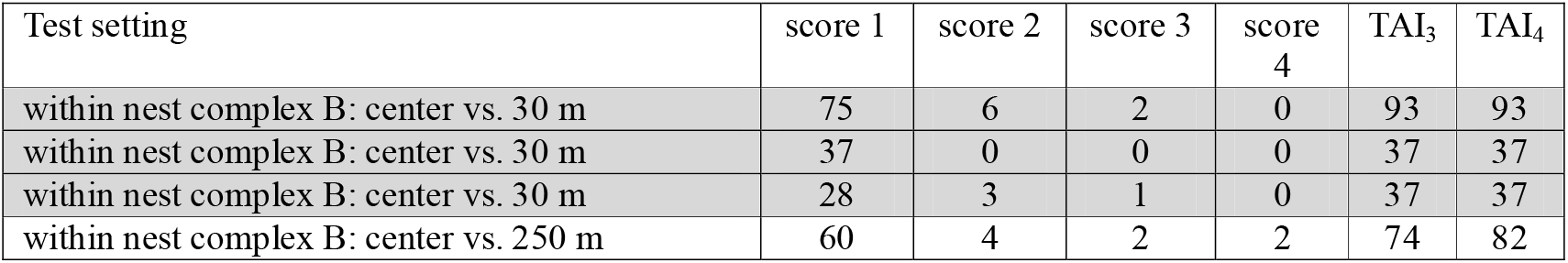
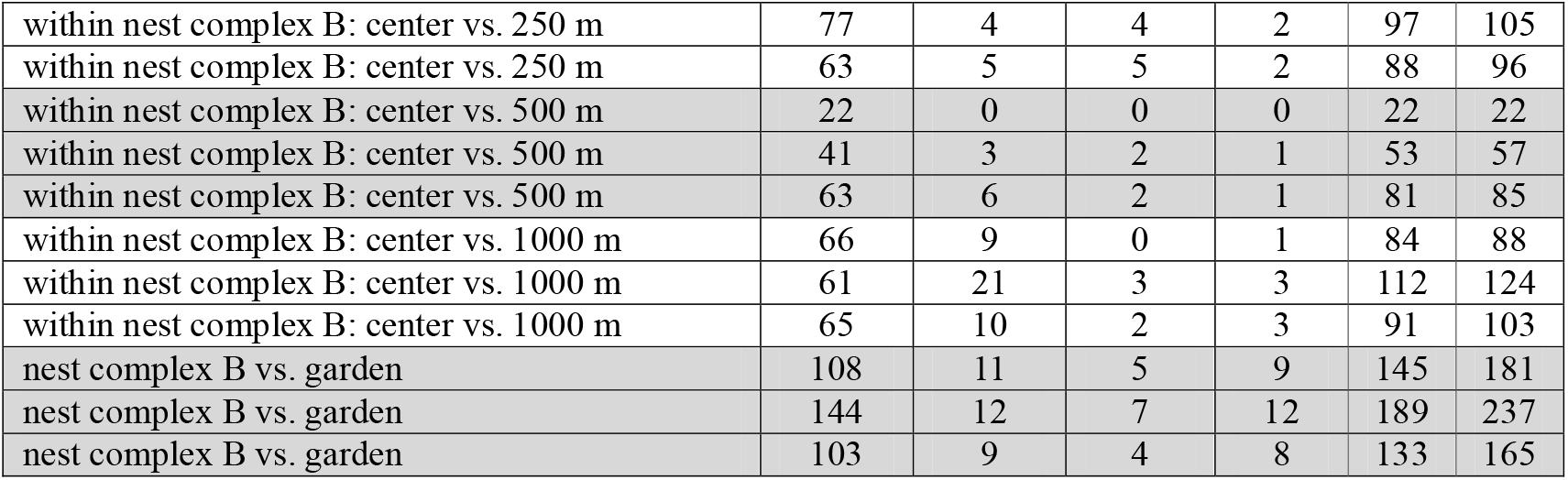
Worker aggression tests between nests from the center of nest complex B and from nests of nest complex B located 30, 250, 500 and 1000 m away from the center (lines 1–12) and aggression tests between nest complex B nests and nests from a remote monodomous garden population (lines 13–15). Increasing scores indicate increasingly aggressive behavior. The Total Agonistic Index TAI_3_ considers only aggressive behavior (scores 1–3) and TAI_4_ additionally the number of killed ants (score 4). Hundred workers from each nest were confronted.

If tested in a one-way ANOVA, the aggression observed between nests within nest complex B did not significantly differ in each of the six pairwise comparisons of the four different test settings. The biggest within-nest complex difference, that of the test setting “center vs. 500 m” and that of the setting “center vs. 1000 m” was also not significant (TAI_3_: F_1,4_ 5.28, p=0.083; TAI_4_: F_1,4_ 5.74, p=0.075). Furthermore, no increase of within-nest complex aggression with growing distance of the nests could be shown in linear regressions of TAI_3_ (r^2^=0.167, n=12, p=0.187) and TAI_4_ data (r^2^=0.203, n=12, p=0.142).

Aggression between nest complex nests and those of the monodomous garden population was significantly larger than in any within-nest complex test. Even the closest data, that of the setting “nest complex vs. garden” and that of the setting “center vs. 1000 m” differed significantly (TAI_3_: F_1,4_ 9.98, p=0.034; TAI_4_: F_1,4_ 13.63, p=0.021). These data give a clear indication that worker aggression within the nest complex is reduced whereas increased agonistic behavior occurs in confrontations of nest complex workers with those from the monodomous (supposedly monogynous) population.

Our observations of moderate aggression in within-nest complex confrontations is suspected to be a consequence of the experimental setting. The encapsulated confrontation within the small space of two tubes suddenly put together with their open ends is likely to raise some initial aggression which would possibly not occur under other conditions. Each ant student who has divided a group of workers in the laboratory, has kept them in separate vials for some days and reunified these by releasing them suddenly into a petri dish will observe initial disorientation accompanied by aggressive behavior which will settle after a while. The same occurs in the field when for instance a *Formica rufa* group nest is massively disturbed and the excited workers attack each moving object including their own nest-mates. The possible boosting of aggression data in our test system does not question our conclusion of a significantly reduced aggression within the nest complex as each test was affected in the same way.

## Discussion

In fact, intensive investigations of the relations between nests on study plots with high nest density of *Lasius niger* have been missing up to the present. What we only have are casual observations of aggressive interactions between nests. According to Czechowski (1984) and Seifert (2018), these typically start with tournaments at territorial borders. Seifert (2018) wrote: “Conflicts between mature colonies of approximately the same size are mainly conducted by ritualized encounters with low mortality: Hundreds of opponents stand face to face along a front line and perform jerky, high-frequent back and forth movements towards the alien ants. This may last one hour or more and is apparently a means of assessing the strength of the opponent. Only in case of strong numeric superiority of one party, the encounter may finally go over to mortal fighting and raiding of the weaker nest.” It is obvious in these cases that the workers of the engaged parties had a disparate pattern of cuticular odor cues and that they most probably belonged to monogynous nests.

Humans are alerted by aggressive acts and not by peace. Hence it is an expected psychological phenomenon that aggressive encounters between ant societies attract the attention of ant students much more than workers peacefully running along trails. This might explain that supercoloniality in *Lasius niger* escaped the perception of observers so far. The ecological dominance and biomass of this ant across its Palaearctic range from Western Europe to Siberia is enormous. The situation in Central Europe was examined by Seifert (2017b). Among 79 study plots on which *Lasius niger* was present, this ant showed densities > 30 nests / 100 m^2^ on ten plots, with a maximum of 108 nests / 100 m^2^ found on a 32 m^2^ plot. He reported the highest densities for grasslands in landscapes with strong anthropogenic impact – often in rural and suburban regions with fertile and moderately dry tschernosem or alluvial soils. In none of these high-density plots, the colony status was checked. Extreme local frequency is also observed in similar habitats in the Ukraine in the Kyiv, Zhytomyr and Chernihiv regions where large associations with hundreds of nest mounds are ubiquitous in rural fields, gardens or initial states of tree plantations. The high abundance of *L. niger* within the city zone of the urban gradient is well known for a long time – for Russia and the Ukraine this has been recently reported for Moscow (Putiatina 2011) and Kyiv (Stukalyuk 2017). Considering these data, we hypothesize that supercoloniality in *Lasius niger* is not the very exception but we also emphasize that the presence of high-density clusters of nests alone is no conclusive argument for the presence of a nest complex. We encourage future ant observers to pay attention to the phenomenon and to collect really indicative data.

## Supporting information

https://docs.google.com/spreadsheets/d/120NPIgE0uSWaq72Ozm4iaQQUekJjyl8H/edit?usp=sharing&ouid=104020205009517280231&rtpof=true&sd=true

## Acknowledgements and declaration of individual contributions

S. Stukalyuk is grateful to W. Czechowski /Warsaw and K. Vepsalainen /Helsinki for help with a primordial version of the manuscript and for suggesting to conduct pleometrosis experiments and to A. Radchenko / Kyiv for determining the ants of the nest complex in the initial phase of the preparation of the paper. We wish to thank M. Kozyr for the botanical investigation in the area of the nest complexes and Galina Stukalyuk for considerable help in collecting material in the field and in maintainment of the pleometrosis experiments. The research leading to this publication has received funding from “Funding the development of priority research fields of the Ukraine” allowance number KPKVK 6541230.

